# Automatic sleep scoring for real-time monitoring and stimulation in individuals with and without sleep apnea

**DOI:** 10.1101/2024.06.12.597764

**Authors:** Martín Esparza-Iaizzo, María Sierra-Torralba, Jens G Klinzing, Javier Minguez, Luis Montesano, Eduardo López-Larraz

## Abstract

Digital therapeutics, enabled by advanced machine learning algorithms and medical wearable devices, offer a promising approach to streamline diagnostics and improve access to healthcare. Within this framework, automatic sleep scoring can provide accurate and efficient sleep analysis from electrophysiological signals recorded with wearable sensors, such as electroencephalography (EEG). However, the optimal configuration and temporal dynamics of automatic sleep scoring systems remain unclear, especially concerning their performance across different population samples. This study systematically investigates the impact of electrode setup, temporal scope, and population characteristics on the performance of automatic sleep scoring algorithms. Utilizing a convolutional neural network (CNN) model, we analyzed various electrode configurations and temporal dynamics using datasets comprising both healthy participants and individuals with sleep apnea. Our findings reveal that sleep scoring based on a single frontal EEG channel demonstrates reliable congruency with human expert scorers, with minimal improvement observed with additional sensors. Moreover, we demonstrate that real-time sleep scoring can be achieved with comparable accuracy to offline methods, which rely on past and future information to classify a window of interest. Remarkably, a notable reduction in decoding accuracy is observed for individuals with sleep disorders compared to healthy participants, highlighting the challenges inherent in accurately assessing sleep stages in clinical populations. Digital solutions for automatic sleep scoring hold promise for facilitating timely diagnoses and personalized treatment plans, with applications extending beyond sleep analysis to include closed-loop neurostimulation interventions. Our findings provide valuable insights into the complexities of automatic sleep scoring and offer considerations for the development of effective and efficient sleep assessment tools in both clinical and research settings.

## Introduction

The rise of digital therapeutics is transforming healthcare. Utilizing advanced software and artificial intelligence (AI) solutions, along with wearable sensors, it is now possible to provide clinicians with tools to monitor or even treat more and more health conditions (Makin 2019; Abbadessa et al. 2022). Sleep medicine is one field that could potentially benefit greatly from this advancement (Park et al. 2021). Disorders of sleep present a significant personal, societal, and economic burden (Sivertsen et al. 2011; Tarasiuk and Reuveni 2013) and are becoming increasingly prevalent (Van Ryswyk et al. 2018; Pavlova and Latreille 2019). Timely sleep diagnostics are crucial for helping individuals with sleep disorders. Unfortunately, waiting times in sleep medicine are very long, especially for diagnostics that involve the recording and analysis of brain activity (Escourrou et al. 2000). Currently, sleep diagnostics rely in large parts on full-night polysomnography (PSG) recordings and manual scoring according to the criteria developed by the American Academy of Sleep Medicine (AASM) (Berry et al. 2015). PSG recordings are evaluated in epochs of 30 seconds, which are labelled with one of five wake/sleep stages: wake (W), REM sleep (R), and the progressively deepening non-REM stages N1-N3. This labelling process is known as *sleep scoring* or *sleep staging*, while the resulting successive representation of all labelled epochs is termed *hypnogram*. Sleep scoring is a time-consuming and resource-intensive process that creates a bottleneck in diagnosis and treatment, highlighting the need for efficient and accessible alternatives. Moreover, despite being performed by trained professionals, manual scoring is subject to a certain degree of uncertainty, with inter-rater reliability being estimated in the range of 80-85% (Danker-Hopfe et al. 2009; Rosenberg and Van Hout 2013). Automatic sleep scoring using machine learning algorithms presents a promising solution. This approach has the potential to streamline the diagnostic process, improve access to sleep healthcare, and ultimately enhance patient outcomes, with applications that could extend beyond sleep analysis, such as closed-loop targeted neurostimulation (Ngo et al. 2013; Lee et al. 2020).

Efforts to develop automatic sleep scoring systems with human-level accuracy began in the late 20th century, utilizing hybrid systems and statistical pattern recognition techniques (Schaltenbrand et al. 1996). Subsequently, various machine learning algorithms, such as decision trees and support vector machines, showed promise in automatic sleep stage classification (Imtiaz and Rodriguez-Villegas 2014; Zhang et al. 2014). However, the recent surge in deep learning research has revolutionized the field. Deep learning architectures, particularly those featuring convolutional neural networks (CNNs), recurrent neural networks (RNNs), and more recently transformers, have achieved remarkable congruency with human expert scorers. Different approaches are commonly compared by their performance on publicly available sleep scoring datasets. Table 1 provides an overview of recently published results, specifically highlighting the most prevalent network architectures employed, but also evidencing significant heterogeneity in other methodological aspects. Firstly, the setup, specifically the number and type of sensors employed for scoring, varies from single electroencephalography (EEG) channels to comprehensive montages, including EEG, electrooculography (EOG), and electromyography (EMG). Secondly, approaches differ in their temporal scope, i.e., whether they utilize past and future information for scoring an epoch. Those that do are limited to offline analyses, while approaches that rely solely on present and past data are suitable for real-time sleep scoring. Thirdly, the sample datasets used for evaluation can be a source of variability. Most studies evaluate different approaches on datasets of healthy participants only, neglecting crucial information on how they perform when analyzing data from patients with sleep disorders.

**Table 1:**
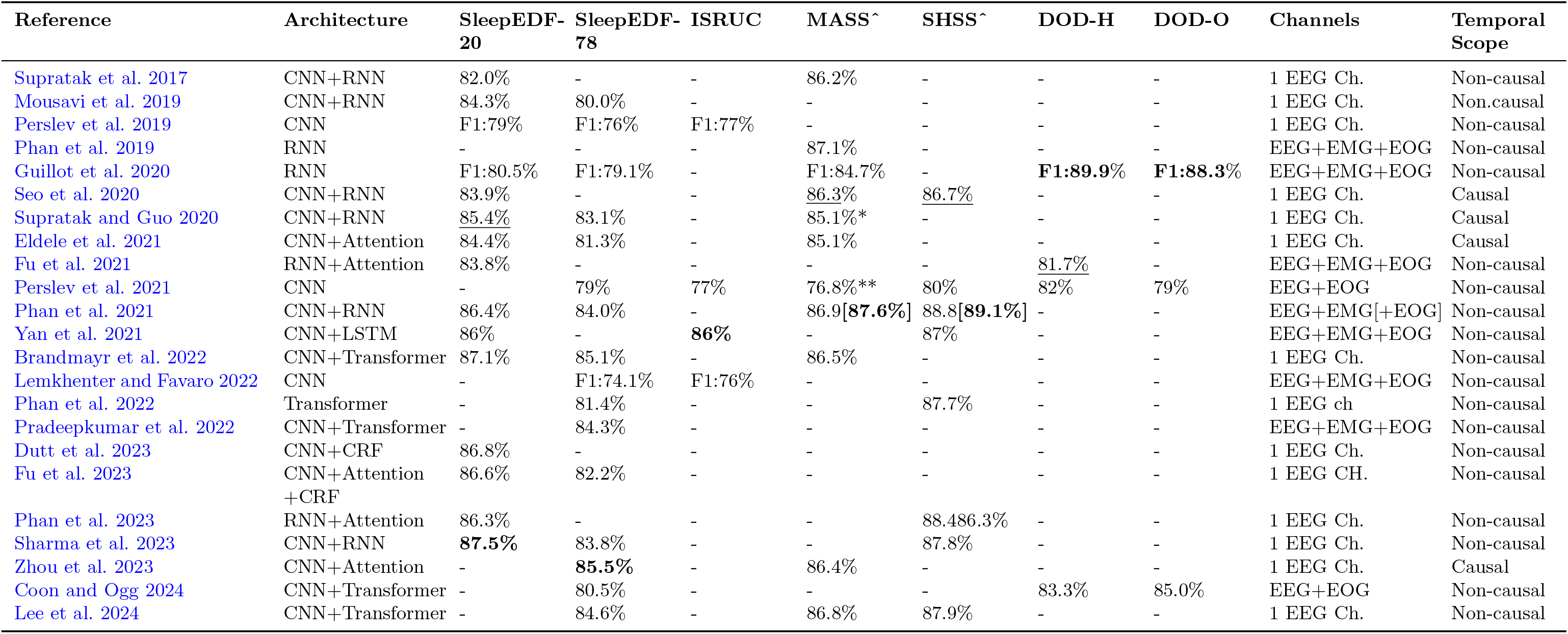
Comparison of recent studies on automatic sleep scoring using deep neural networks. The table presents reported performance metrics from various studies on their respective datasets. Decoding accuracy results are provided for most studies except for those that only report the F1 score (marked as F1). For studies comparing different methodologies, such as different electrode configurations, the highest decoding accuracy is reported. The best result for each dataset is marked in bold. The best result obtained with purely online methods is underlined. CNN: convolutional neural network; RNN: recurrent neural network; CRF: conditional random fields; SleepEDF-20: 2013 version of the SleepEDF dataset, including 20 participants; SleepEDF-78: 2018 version of the SleepEDF dataset, including 78 participants (Kemp et al. 2000); ISRUC: ISRUC-Sleep Dataset (Khalighi et al. 2016); MASS: Montreal Archive of Sleep Studies (O’Reilly et al. 2014); SHSS: Sleep Heart Health Study (Quan et al. 1997); DOD-H: Dreem Open Dataset-Healthy; DOD-O: Dreem Open Dataset-Obstructive (Guillot et al. 2020). * value obtained as the weighted average of the percentages reported for each MASS subset; ** value obtained as the weighted average of the percentages reported for MASS subsets 1 and 3. approval required to access these datasets.

| Reference | Architecture | SleepEDF-20 | SleepEDF-78 | ISRUC | MASS<sup>^</sup> | SHSS<sup>^</sup> | DOD-H | DOD-O | Channels | Temporal Scope |
| --- | --- | --- | --- | --- | --- | --- | --- | --- | --- | --- |
| Supratak et al. 2017 | CNN+RNN | 82.0% | - | - | 86.2% | - | - | - | 1 EEG Ch. | Non-causal |
| Mousavi et al. 2019 | CNN+RNN | 84.3% | 80.0% | - | - | - | - | - | 1 EEG Ch. | Non-causal |
| Perslev et al. 2019 | CNN | F1:79% | F1:76% | F1:77% | - | - | - | - | 1 EEG Ch. | Non-causal |
| Phan et al. 2019 | RNN | - | - | - | 87.1% | - | - | - | EEG+EMG+EOG | Non-causal |
| Guillot et al. 2020 | RNN | F1:80.5% | F1:79.1% | - | F1:84.7% | - | **F1:89.9%** | **F1:88.3%** | EEG+EMG+EOG | Non-causal |
| Seo et al. 2020 | CNN+RNN | 83.9% | - | - | 86.3% | <u>86.7%</u> | - | - | 1 EEG Ch. | Causal |
| Supratak and Guo 2020 | CNN+RNN | <u>85.4%</u> | 83.1% | - | 85.1%* | - | - | - | 1 EEG Ch. | Causal |
| Eldele et al. 2021 | CNN+Attention | 84.4% | 81.3% | - | 85.1% | - | - | - | 1 EEG Ch. | Causal |
| Fu et al. 2021 | RNN+Attention | 83.8% | - | - | - | - | <u>81.7%</u> | - | EEG+EMG+EOG | Non-causal |
| Perslev et al. 2021 | CNN | - | 79% | 77% | 76.8%** | 80% | 82% | 79% | EEG+EOG | Non-causal |
| Phan et al. 2021 | CNN+RNN | 86.4% | 84.0% | - | 86.9[**87.6%**] | 88.8[**89.1%**] | - | - | EEG+EMG[+EOG] | Non-causal |
| Yan et al. 2021 | CNN+LSTM | 86% | - | **86%** | - | 87% | - | - | EEG+EMG+EOG | Non-causal |
| Brandmayr et al. 2022 | CNN+Transformer | 87.1% | 85.1% | - | 86.5% | - | - | - | 1 EEG Ch. | Non-causal |
| Lemkhenter and Favaro 2022 | CNN | - | F1:74.1% | F1:76% | - | - | - | - | EEG+EMG+EOG | Non-causal |
| Phan et al. 2022 | Transformer | - | 81.4% | - | - | 87.7% | - | - | 1 EEG ch | Non-causal |
| Pradeepkumar et al. 2022 | CNN+Transformer | - | 84.3% | - | - | - | - | - | EEG+EMG+EOG | Non-causal |
| Dutt et al. 2023 | CNN+CRF | 86.8% | - | - | - | - | - | - | 1 EEG Ch. | Non-causal |
| Fu et al. 2023 | CNN+Attention<br>+CRF | 86.6% | 82.2% | - | - | - | - | - | 1 EEG CH. | Non-causal |
| Phan et al. 2023 | RNN+Attention | 86.3% | - | - | - | 88.486.3% | - | - | 1 EEG Ch. | Non-causal |
| Sharma et al. 2023 | CNN+RNN | **87.5%** | 83.8% | - | - | 87.8% | - | - | 1 EEG Ch. | Non-causal |
| Zhou et al. 2023 | CNN+Attention | - | **85.5%** | - | 86.4% | - | - | - | 1 EEG Ch. | Causal |
| Coon and Ogg 2024 | CNN+Transformer | - | 80.5% | - | - | - | 83.3% | 85.0% | EEG+EOG | Non-causal |
| Lee et al. 2024 | CNN+Transformer | - | 84.6% | - | 86.8% | 87.9% | - | - | 1 EEG Ch. | Non-causal |

The present paper addresses these three aspects (setup, scope, and sample) by systematically analyzing their impact on automatic sleep scoring. Using a previously developed CNN model (Youness 2020), we compared the model’s performance using different combinations of physiological signals, ranging from single EEG derivations on different head locations, to multimodal polysomnography recordings. Additionally, we investigated the impact of temporal dynamics by analyzing performance when considering only present and past data, or different amounts of future data. The evaluations are conducted on four popular open-access datasets, two of which include only healthy participants, and the other two with participants with sleep disorders. By investigating these factors, we aim to identify the optimal channel configuration and temporal scope for accurate sleep staging in real-time applications. Our results improve the understanding of automatic sleep scoring for diagnostics and lay the groundwork for its use in therapeutic sleep interventions, such as closed-loop neurostimulation.

## Methods

To evaluate the performance of our automatic sleep scoring model under diverse conditions and assess its generalizability across different populations, we utilized four open-access datasets. These datasets encompass recordings from both healthy individuals and patients diagnosed with sleep disorders, allowing for a comprehensive assessment of the model’s robustness. Furthermore, the datasets include a variety of sensor configurations, comprising EEG, EOG, EMG, and other physiological signals. This variability enabled us to explore the impact of sensor selection on sleep staging accuracy and to investigate the potential for real-time applications with limited sensor setups. Table 2 provides an overview of the key characteristics of each dataset, including demographic information, the number of recordings, available physiological signals, and sampling rate.

**Table 2:** Summary of datasets used in the study. The four datasets comprise recordings from both healthy participants and patients diagnosed with sleep apnea. All datasets contained multiple EEG channels as well as other biosensors.

| Dataset | Population | Files | Subjects<br>(M/F) | Age<br>(yrs) | EEG Ch. | EEG<br>Rate | Additional<br>sensors |
| --- | --- | --- | --- | --- | --- | --- | --- |
| SleepEDF | Healthy | 153 | 78<br>(37/41) | 58.8±22.1 | Fpz-Cz, Pz-Oz | 100 | EOG, EMG,<br>oro-nasal and<br>rectal tempera-<br>ture |
| ISRUC | Sleep apnea | 100 | 100<br>(55/45) | 51±16 | F4-A1, C4-A1,<br>O2-A1, F3-A2,<br>C3-A2, O1-A2 | 200 | EOG, EMG,<br>ECG, Snore,<br>pressure-based<br>airflow, SaO2<br>and body posi-<br>tion |
| DOD-H | Healthy | 25 | 25 (19/6) | 35.3±7.5 | C3-M2, F4-M1,<br>F3-F4, F3-M2,<br>F4-O2, F3-O1,<br>FP1-F3, FP1-M2, FP1-O1,<br>FP2-F4, FP2-M1, FP2-O2 | 250 | EOG, EMG,<br>and ECG |
| DOD-O | Sleep apnea | 55 | 55<br>(35/20) | 45.6±16.5 | C3-M2, C4-M1,<br>F3-F4, F3-M2,<br>F4-O2, F3-O1,<br>O1-M2, O2-M1 | 250 | EOG, EMG,<br>and ECG |

In detail, the SleepEDF Database Expanded (Goldberger et al. 2000; Kemp et al. 2000) is an open-access dataset comprised of 197 20-hour PSG recordings with EEG, EOG, EMG, respiration and body temperature data. The data is divided into 2 cohorts, the Sleep Telemetry Study and the Sleep Cassette Study, formed by patients under the effects of temazepam and control volunteers, respectively. Analyses on this dataset have been conducted on its 2013 version (SleepEDF-20), which included data from 20 participants, and on its 2018 “Expanded” version (SleepEDF-78), which includes data from 78 participants. In this study, we utilized the Sleep Cassette cohort of the SleepEDF-78 dataset, consisting of 153 recordings from 78 participants. Available EEG channels are sampled at 100 Hz and were recorded from Fpz–Cz and Pz–Oz locations. An initial set of preprocessing steps was implemented as suggested by (Supratak et al., 2017). First, stages marked as movement (M) or not scored (NS) were removed. Secondly, stages S3 and S4 were merged into N3 to ensure consistency with the AASM scoring system. Finally, each night was extracted from the 20-hour PSG by cropping the recording 30 minutes before and after sleep onset/offset.

The ISRUC-Sleep dataset (Khalighi et al. 2016) is composed of three subgroups (I-III) of participants scored by two human experts according to the AASM guidelines. Data from subgroups I and II include 100 and 8 patients, respectively, with evidence of sleep apnea events, while subgroup III comprises 10 healthy controls. Given the large imbalance in the number of subjects, subgroup I was exclusively employed. PSG recordings contain information from various sensors including EOG, EEG (F3–A2, F4–A1, C3–A2, C4–A1, O1–A2, and O2–A2), EMG, ECG, SaO2, position, airflow, and abdominal effort. Electrical activity is sampled at 200 Hz and EEG signals are band-passed filtered between 0.3 and 35 Hz and notch-filtered at 50 Hz.

Finally, we utilized the two Dreem Open Datasets (Guillot et al. 2020), including healthy volunteers and patients diagnosed with obstructive sleep apnea (OSA), respectively. The first dataset, DOD-H (Dreem Open Dataset – Healthy), contains 25 PSG recordings scored by 5 expert sleep technologists following AASM criteria. Available data include EOG, EMG, ECG, and 12 EEG channels (C3–M2, F4–M1, F3–F4, F3–M2, F4–O2, F3–O1, FP1–F3, FP1–M2, FP1–O1, FP2–F4, FP2–M1, FP2–O2) sampled at 250 Hz. The second dataset, DOD-O (Dreem Open Dataset – Obstructive), consists of 55 PSG recordings from a cohort of patients and is labeled by 5 experienced scorers as in DOD-H. The available data differs only in the number of EEG channels, which drops from 12 to 8 (C3–M2, C4–M1, F3–F4, F3–M2, F4–O2, F3–O1, O1–M2, O2–M1), also sampled at 250 Hz.

## Model

### Preprocessing

The general preprocessing pipeline is depicted in Figure 1A. Every PSG recording contains a variable number of channels sampled at different frequencies and divided into a discrete number of labels associated with each epoch. Initially, each file is resampled to 100 Hz and reshaped into a matrix where the first dimension is the number of epochs, the second is the number of time points (3000), and the third is the employed number of channels, which depends on the test being carried out. The subjects in each dataset are then divided into training and testing with a 90% and 10% ratio, respectively, to carry out a 10-fold cross-validation. The training data is furthermore partitioned into a train (85%) and validation set (15%). Finally, the recordings are z-scored exclusively according to the training data, ensuring that we do not employ information from the test set in any way.

**Figure 1:**
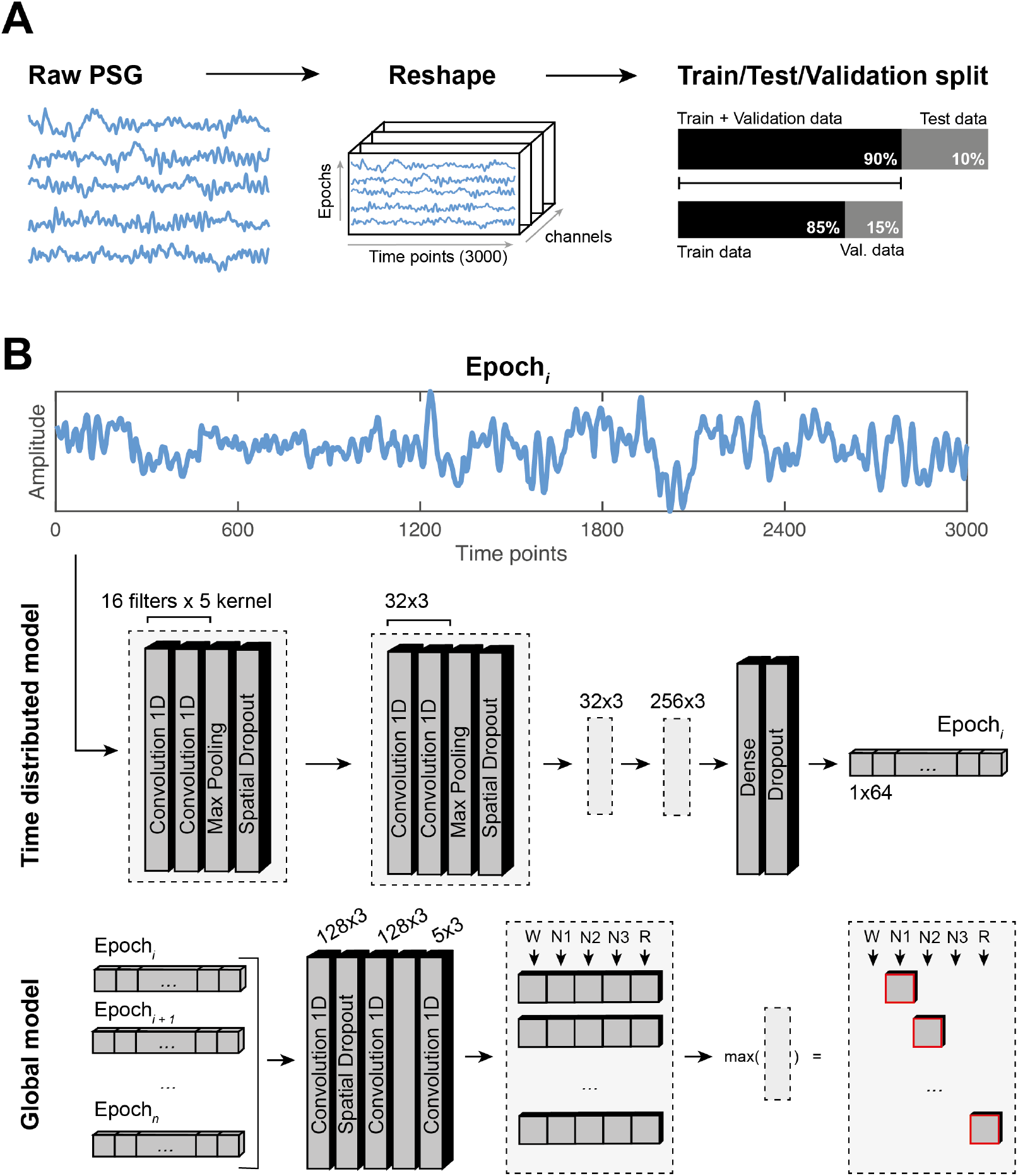
Preprocessing and CNN model. (A) Preprocessing pipeline. PSG recordings are resampled and reshaped into a 3-dimensional matrix representing epochs, time points, and channels. Subsequently, the data is divided into training, validation, and test sets. (B) CNN architecture. The model comprises two submodels. Firstly, the ‘Time Distributed’ step encodes each epoch independently of their sequential order. Then, the resulting epoch-encoded vectors (1x64) are fed into the ‘Global’ submodel, which generates the final output labels sequentially.

### Architecture

The convolutional neural network (CNN) used as the basis of comparison in this study was originally proposed in (Youness 2020). From a broad perspective, the architecture comprises two main submodels (see Figure 1). The first, referred to as the ‘Time Distributed’ submodel, is responsible for feature extraction of each individual epoch, irrespective of its position during the night and without considering neighboring information. The second submodel, referred to as the ‘Global’ submodel, takes N epochs encoded by the Time Distributed submodel and processes them to derive the corresponding sleep scores.

In the Time Distributed submodel, data points of each epoch pass through a structure of two 1D convolutions of 16 filters and a kernel size of 5. Afterwards, pooling of size 2 and spatial dropout with rate 0.01 is performed. This structure (1D Conv, 1D Conv, pooling, and dropout) is repeated 4 times with an increasing number of filters from 16 to 32 (twice) and finally 256, while reducing the kernel size to 3 in all cases. Finally, a fully connected dense layer of 64 values is created and used as output for every epoch.

The Global model takes the 64-unit vectors and performs another series of operations on them. This implies that weights are no longer applied independently to each epoch, but rather to the entirety of the training, validation, or testing batch. These operations comprise two sets of 1D convolutions (128 filters and a kernel size of 3) and spatial dropout with a rate of 0.01, followed by a final 1D convolution (5 filter and a kernel size of 3) with softmax activation to obtain a 5-unit (W, N1, N2, N3 and R) probabilistic vector. The optimization algorithm selected is Adam (Learning rate = 0.0001) (Kingma and Ba 2014), and the loss function employed is the sparse categorical cross-entropy since labels are not one-hot encoded. All activation functions except for the final layer are rectified linear units (ReLU).

### Training and inference

In both the causal and non-causal scenarios, PSG data were fed in batches of L=100 epochs during training, to accelerate the convergence of the model and reduce processing time. All output labels were considered to compute the loss and backpropagate the errors. Nevertheless, despite the dimension of L, the kernel size of 3 used in the convolutional layers of the sequence learning part directly influences the receptive field of the neurons during the convolution operation. Thus, when using a non-causal configuration, the information captured during training is derived from the previous, the current, and the subsequent epochs. When it comes to the causal configuration, padding was applied to the rightmost epoch, to prevent the network from accessing future features during training. The same can be extrapolated to inference, although in this regard there are more differences between the two scenarios described.

For causal inference (i.e., real-time compatible), we employed a sliding window with a batch size reduced to L=5 epochs, corresponding to 2.5 minutes. This minimized the size of the time window required for testing, a paramount trait of online systems. Furthermore, the stride of the sliding time window was set to 1, and for every new batch of 5 epochs, we only retained the final label, since our aim was to mimic real-time signal acquisition. Conversely, for non-causal inference (i.e., only usable for offline applications), we used a batch size of 100 non-overlapping epochs and kept all predicted labels. Accordingly, batching remained identical to training in this case.

The number of channels employed during training and testing differed according to the analysis being carried out (see 2.3. Systematic Analysis). For example, for our control scenario, the number of channels was set to 1, as training and inference were done on single-channel EEG data. However, for the multi-channel analysis, the number of channels increased depending on the employed electrophysiological data.

The model was implemented in Python programming language v3.7, with Keras API v2.3.0 running TensorFlow v2.5.0 as the backend (Van Rossum and Drake 1995; Abadi et al. 2016; Chollet 2018). To maximize the available datasets and assess the generalization capabilities of the network, we performed a 10-fold cross-validation at every analysis stage. Training, validation, and testing are carried out on a GPU NVIDIA GeForce RTX 2060 with sequences of size L=100 being processed together in each training batch, 100 epochs in each training fold. 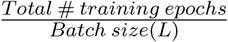 steps per training epoch, and 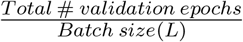 validation steps taken at the end of each training epoch.

### Systematic analysis

We conducted a comparison of this model in a series of different configurations. Our control condition was a single-channel EEG-based causal configuration, which, a priori, provides the most limited information for decoding but offers significant potential in clinical and research settings (Andreotti et al. 2018). The systematic analysis aims to answer four questions regarding the impact of the following aspects on scoring accuracy: i) EEG electrode location, ii) EEG electrode number, iii) incorporation of multimodal data, and iv) available past and future information during decoding (see Figure 2). Additionally, we analyzed the accuracy differences obtained for healthy participants compared to sleep apnea patients.

**Figure 2:**
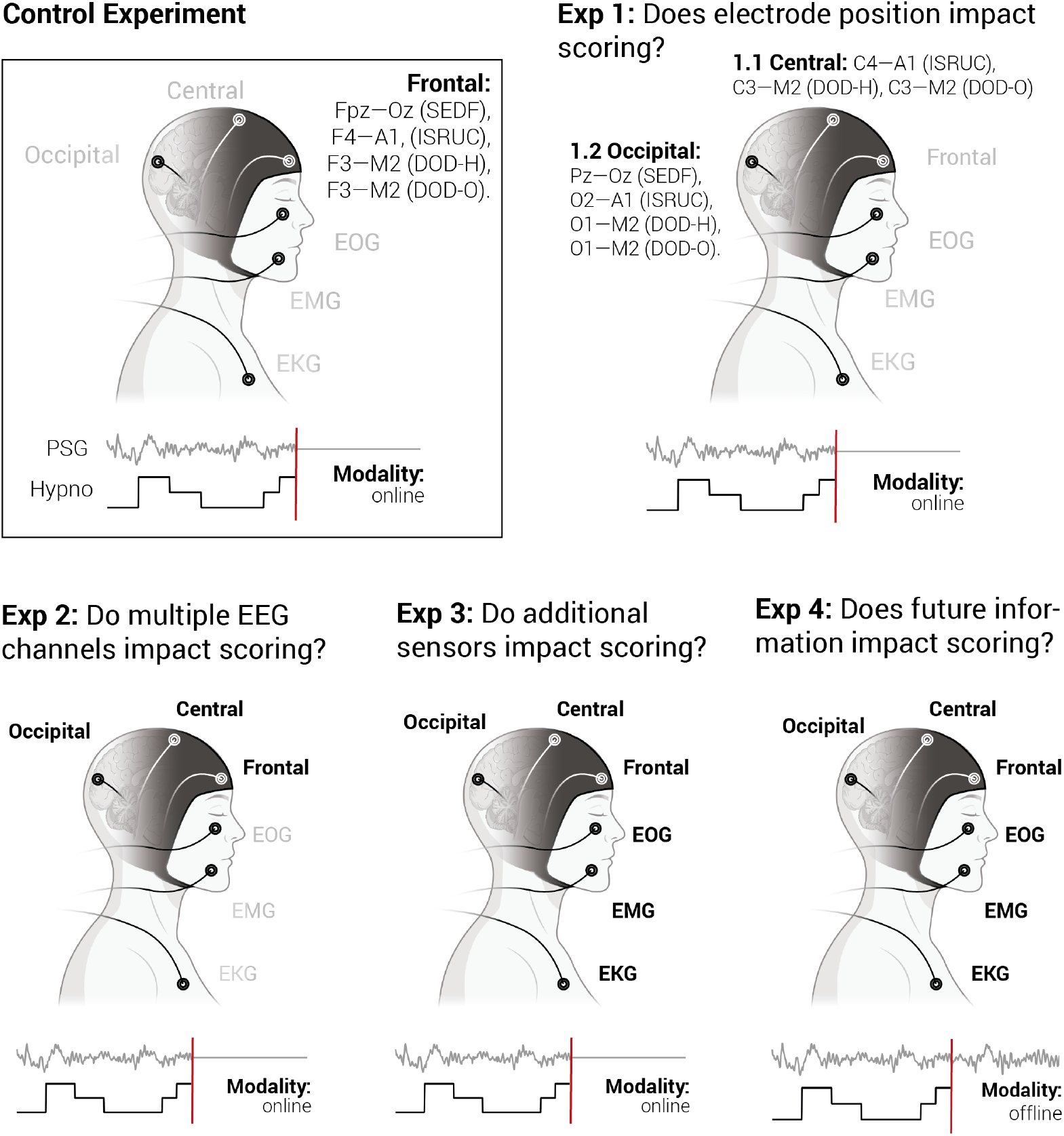
Systematic analysis flow chart. (A). The series of four experiments aims at answering four questions regarding electrode positioning, the relevance of using multiple EEG contacts, the use multimodal data, and the influence of available future information. The employed derivations and staging modality (causal/non-causal) are presented for each experiment. The control scenario employed a single-channel EEG-based real-time setup using the same CNN architecture (see Figure 1).

The control condition employed the following derivations: [Fpz–Cz] in SleepEDF; [F3–A2] in ISRUC; [F3–M2] in DOD-H; and [F3–M2] in DOD-O. The specific location of each input channel (i.e., Fpz vs. F3) depended on the availability within the original dataset. Whenever possible, we employed the same derivation to be consistent in our analysis. As aforementioned, this condition encompassed a single frontal channel across datasets coupled with causal inference using a 2.5-minute sliding time window. Furthermore, we employed causal inference for all experiments except the final one, which precisely aimed to compare this aspect with non-causal testing.

Experiment 1 compared different EEG electrode locations to the control scenario. The underlying question was to understand which recording sites are optimal for staging or if, contrarily, relevant EEG hallmarks (e.g., SOs and spindles) are sufficiently captured from any position. In greater detail, we carried out two sub-experiments at this level, where the staging setup employed single-channel central (Experiment 1.1) and occipital (Experiment 1.2) derivations across all four datasets. The central experiment used [C3–A2] in ISRUC, [C3–M2] in DOD-H, and [C3–M2] in DOD-O; while the occipital experiment employed: [Pz–Oz] in SleepEDF, [O1–A2] in ISRUC, [O1–M2] in DOD-H, and [O1–M2] in DOD-O. Note that the derivation O1–M2 in DOD-H had to be obtained by subtracting channels F3–M2 and F3–O1.

Experiment 2 consisted of evaluating whether a combination of EEG channels from all three locations provided additional information that could improve scoring over a single-channel approach. In this case, the staging configuration combined the previously described EEG derivations from each dataset to create a double-channel input in SleepEDF ([Fpz–Cz, Pz–Oz]), and a triple channel input in ISRUC ([F3–A2, C3–A2, O1–A2]), DOD-H, and DOD-O ([F3–M1, C3–M2, O1–M2] for both). Experiment 3 aimed to take this one step further by assessing if additional information from EOG, EMG and ECG channels could improve the algorithm results. Regarding SleepEDF, the only additional sensor available was chin EMG and therefore, the final configuration was [Fpz–Cz, Pz–Oz, EMG]. For ISRUC, we used [F3–A2, C3–A2, O1–A2, EMG, ECG, EOG]. Finally, in both DOD-H and DOD-O, we employed [F3–M1, C3–M2, O1–M2, EMG, ECG, EOG].

Experiment 4 was designed to assess whether non-causal inference improved classification performance and whether a higher number of neighboring epochs appearing in the context of a target epoch provided advantages in its recognition. Here, we reverted to using a single frontal EEG electrode to limit the number of changing conditions and perform a faithful comparison with our control condition. Note that, in this configuration, data is batched in non-overlapping windows of 100 epochs, whereas all previous stages employed a sliding window of 5 epochs.

### Evaluation metrics

Performance of the model is evaluated with confusion matrices and a series of classification metrics to assess the agreement between the expert’s labels and the algorithm. The first employed metric in the classification report is accuracy,

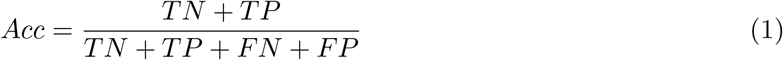

where TP, TN, FP, and FN represent true positives, true negatives, false positives, and false negatives, respectively. Furthermore, precision (or positive predictive value) and recall (or sensitivity) are also reported and defined as:

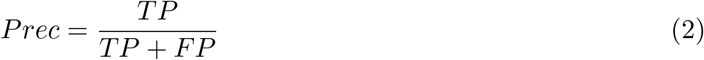

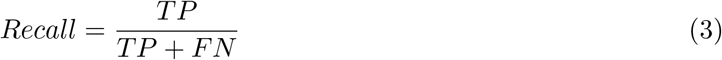

Lastly, we calculate F1 score, which is the harmonic mean of precision and recall:

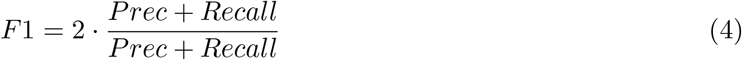

## Results

Table 3 presents an overview of the accuracy metric of these results, which is further segregated by dataset and evaluation experiment. Supplementary Table S1 shows the F1 score following the same structure.

**Table 3:** Accuracy results overview. Decoding accuracy achieved for each dataset in the control scenario and in the different evaluation experiments.

| Accuracy (%) | Control | Exp. 1.1 | Exp.1.2 | Exp. 2 | Exp. 3 | Exp. 4 |
| --- | --- | --- | --- | --- | --- | --- |
| SleepEDF | 80.97 | - | 77.12 | 80.90 | 82.54 | 82.03 |
| DOD-H | 81.92 | 80.79 | 76.99 | 81.56 | 68.00 | 82.91 |
| ISRUC | 71.69 | 75.15 | 65.20 | 71.67 | 76.42 | 76.27 |
| DOD-O | 79.31 | 78.24 | 73.83 | 79.11 | 72.82 | 81.38 |
| **Average** | 78.47 | 78.06 | 73.29 | 78.31 | 74.95 | 80.65 |

The upcoming sections present the results averaged between healthy participants and patients to focus on the comparison, which is oriented towards single-versus multi-channel modalities and causal and non-causal approaches. More specifically, we will present the single-channel (frontal derivation) online staging results in Section 3.1 and employ this as our control scenario. From there onwards, all ensuing results are compared to this decoding configuration as we explore the benefits of introducing additional features into our model. Finally, we recapitulate our findings by dissecting the results between healthy participants and patients to provide further insight on the effects of altered sleep on staging.

### Single-channel online decoding

The results from our control configuration, which employs single channel decoding from frontal derivations across datasets, are presented in Figure 3. Wake, N2, and N3 classes reach the highest values of accuracy, precision, recall, and F1-score, around 80%, while REM (R) falls to approximately 70% in all metrics. The N1 phase presents the lowest results at 37%, with most of the model’s confusion contained within Wake or N2 (see Figure 3A-B). The combination of N2 and N3, furthermore, exhibits almost perfect specificity, with very few examples of either class being wrongly predicted as W, N1, or REM (R). Training and validation accuracies across epochs are shown in Figure 3C, where validation (blue trace) clearly stagnates after approximately 40 epochs. Additionally, Figure 3D presents a hypnogram comparison between the expert’s ground truth and our model’s predictions for a single sample night. Here, we can observe how the confusion from panel A is further present in the night, with a low number of N1 states misclassified and multiple N3 states confused as N2.

**Figure 3:**
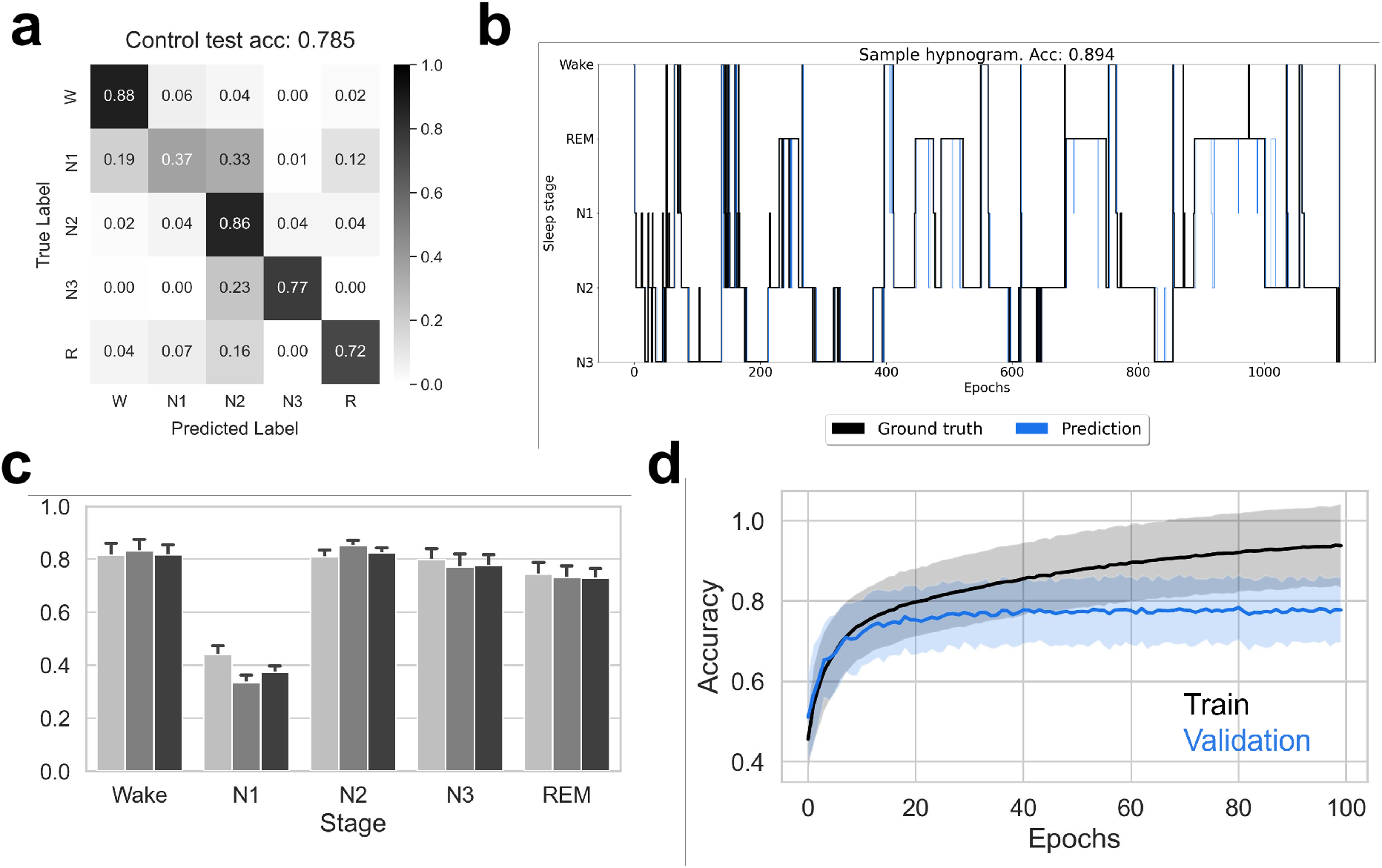
Single channel real-time decoding performance: Control scenario. Panel A shows the confusion matrix together with the average accuracy value obtained in the Control scenario. Panel B contains a bar plot depicting the values of recall (light grey), F1-score (dark grey), and precision (black). Panel C displays the accuracy curve during training (black) and validation (blue). Panel D illustrates the comparison between the ground truth hypnogram annotated by the sleep expert (black) and the one predicted by the CNN model for a sample night’s sleep cycle.

### Comparison of single-channel location for real-time decoding

Understanding the contributions of each electrode to the overall real-time staging accuracy is crucial for discerning the essential information needed to accurately decode sleep stages and fully leverage the potential of single-channel automatic labeling. To address this question, we compared the accuracy of our model across two additional commonly used sites during polysomnographic studies. While the average decoding accuracy for the Control test (frontal electrode) was 78.5%, it dropped to 78.1% and 73.3% for central and occipital electrodes, respectively. Figure 4A details these results and showcases the clear decrease in accuracy when employing occipital derivations. This decline is particularly impactful in the case of N1 and REM scoring, with an 11% and 10% drop in accuracy.

**Figure 4:**
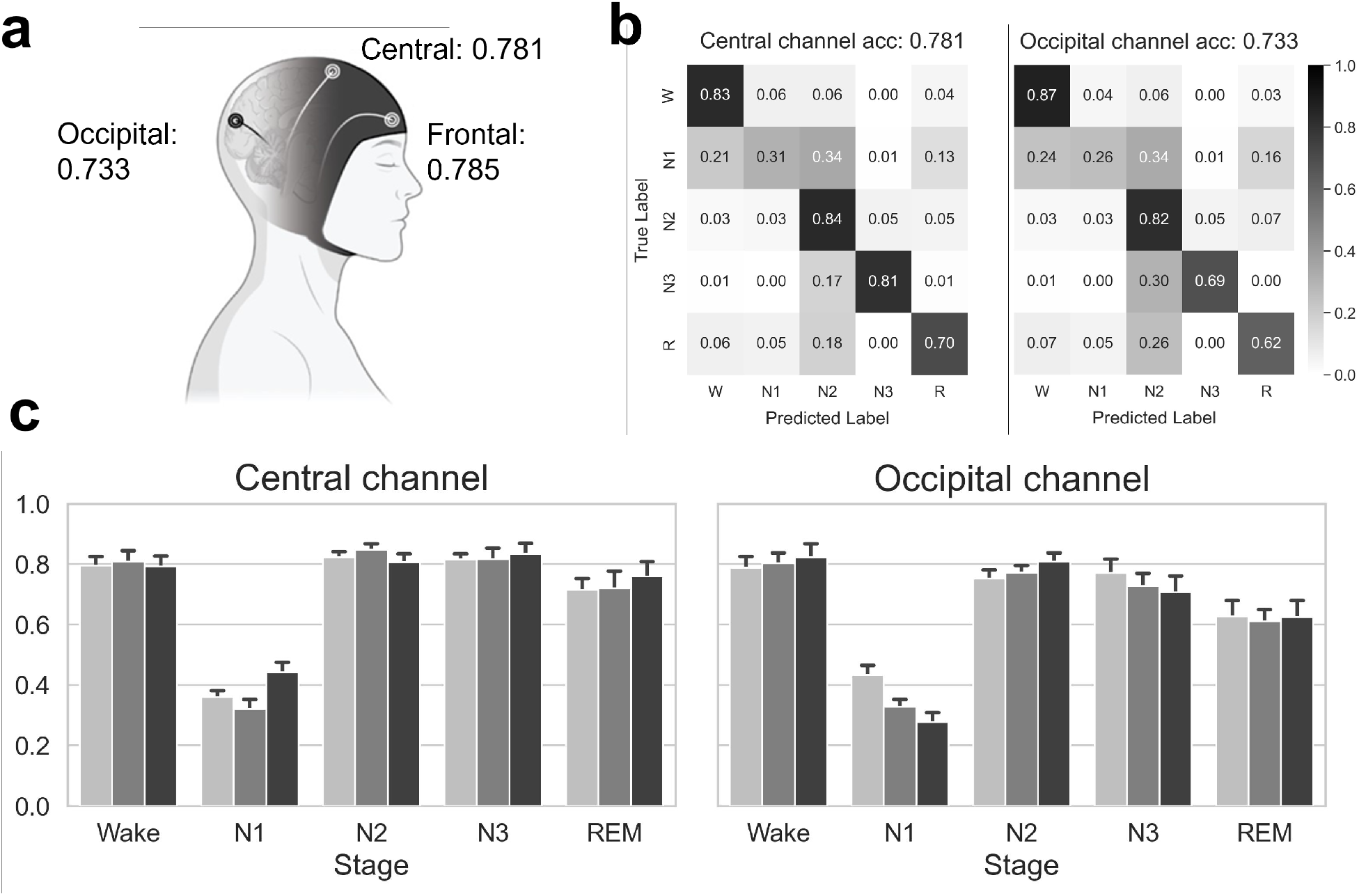
Single channel real-time decoding performance: Experiment 1. Panel A presents an overview of the accuracy values achieved when using information from different electrode positions for decoding (frontal – Control scenario, central – Experiment 1.1, and occipital – Experiment 1.2). Panel B displays the confusion matrices together with the accuracy value obtained in Experiments 1.1 and 1.2. Panel C contains bar plots showing the values of recall (light grey), F1-score (dark grey), and precision (black) for these experiments.

### Multiple-channel real-time decoding

Despite the improvement observed with the use of frontal electrode location compared to central or occipital positions, we aimed to ascertain if each region provided complementary information. This would suggest that adding central and occipital features to frontal derivations could enhance decoding accuracy. Conversely, if frontal regions already encompassed all relevant features present in other locations, one would not expect the results to improve notably. This concept extends to channels beyond EEG, such as EMG and EOG, and ECG, routinely utilized by physicians during manual sleep staging (Figure 5).

**Figure 5:**
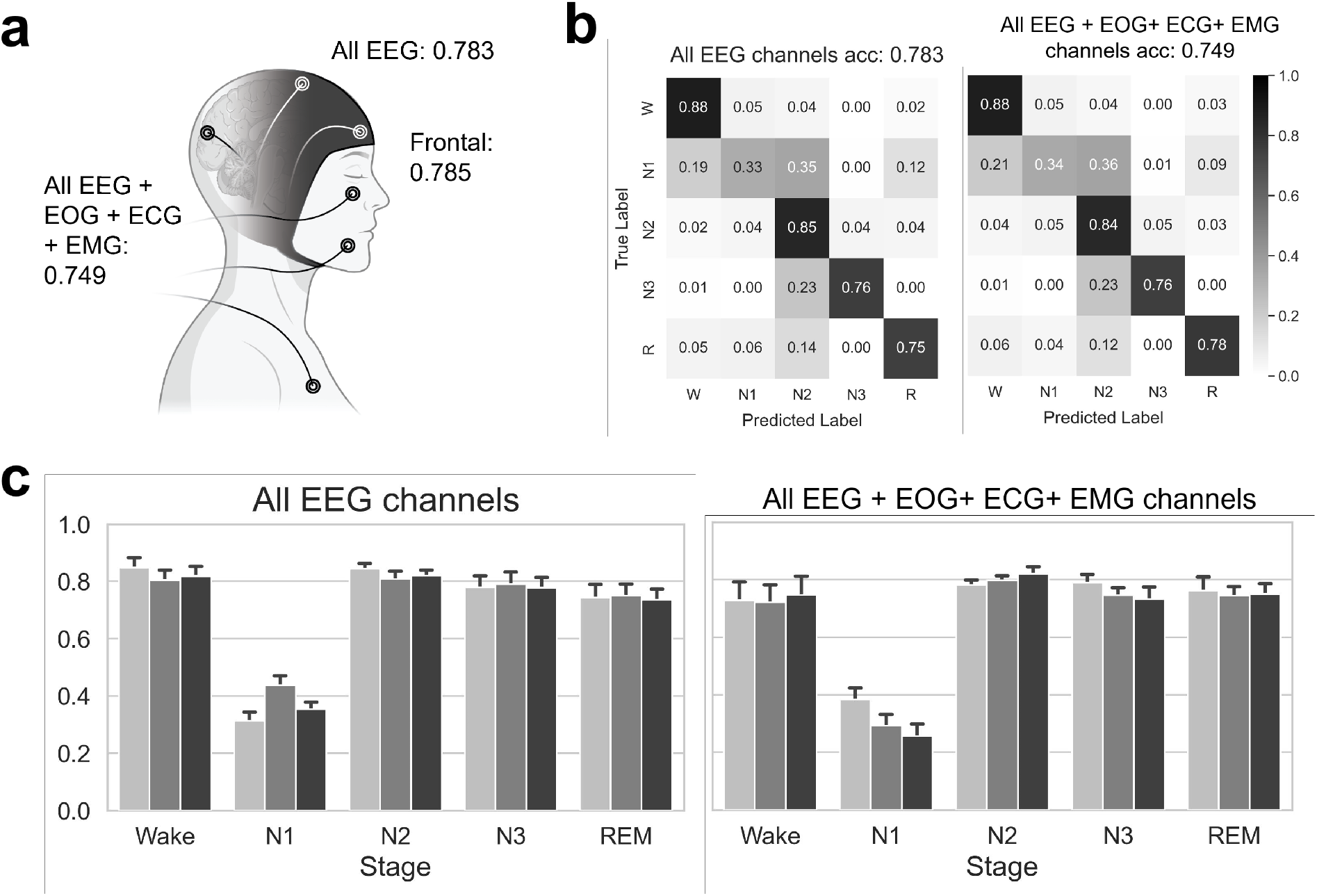
Multiple channel real-time decoding performance: Experiments 2 and 3. Panel A presents an overview of the accuracy values achieved when combining the information from all available EEG derivations, as well as when also incorporating all available channels from different modalities for decoding (all EEG – Experiment 2, all EEG + EOG +ECG + EMG – Experiment 3). Panel B shows the confusion matrices together with the accuracy value obtained in Experiments 2 and 3. Panel C contains bar plots displaying the values of recall (light grey), F1-score (dark grey), and precision (black) for such experiments.

To explore this concept, we combined all available EEG derivations from frontal, central, and occipital locations, observing no significant increase in overall accuracy compared to using frontal locations only (78.3% vs 78.5%, see Figure 5A and B). Extending this analysis to all available electrophysiological signals, including EOG, EMG, ECG, and EEG derivations, not only failed to improve the results, but resulted in a 3.6% decrease compared to the control condition (74.9% vs 78.5%, see Figure 5A and B). These changes in accuracy were similarly reflected in precision, recall and F1 score (Figure 5C).

### Single-channel non-causal decoding

Most automatic sleep staging paradigms utilize both future and past information from single or multiple channels when inferring labels of a specific 30-second EEG/PSG epoch (cf. Table 1). In the context of offline sleep classification, looking at future data to assign a label to an epoch could result in higher performance; however, its main limitation lies in the lack of transferability to real-time staging implementation. In this section, we studied if and how offline approaches, which consider future information to label an epoch, actually outperform our real-time compatible Control experiment (first presented in Figure 6).

**Figure 6:**
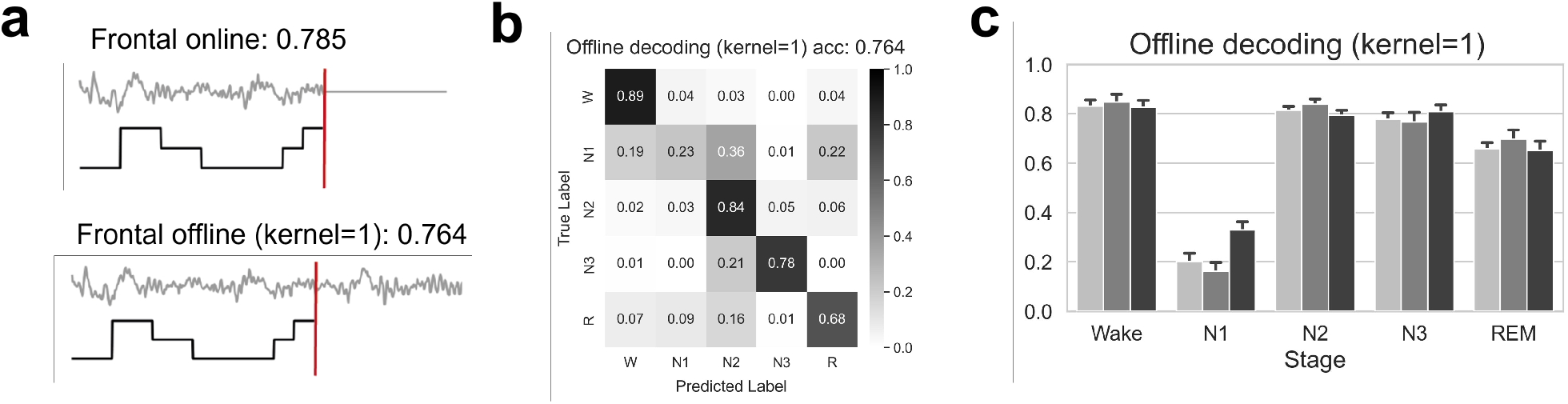
Single-channel non-causal decoding performance: Experiment 4. Panel A presents an overview of the accuracy values achieved in the Control scenario and in its analogue offline approach (kernel = 3, Experiment 4). Panel B shows the confusion matrices together with the accuracy value obtained in Experiment 4. Panel C contains bar plots displaying for such experiment the values of recall (light grey), F1-score (dark grey), and precision (black).

We first analyzed the causal and non-causal implementations of the model as described in the “2.2.3. Training and inference” subsection, with a kernel size of 3 for the sequence learning (which in the case of the online operation, has an effective size of 2, as its rightmost epoch is padded in training and inference). We compare the results from our control scenario to its identical offline counterpart (see Figure 6A). Here, single-channel EEG data are evaluated either in a causal configuration, which, to infer the label of an epoch considers such epoch and its predecessor (i.e., real-time compatible), or in a non-causal manner, which also includes the information of the subsequent epoch. As can be seen in Figure 6, the difference between both approaches is of about 2%.

The number of neighboring epochs in an input sequence influencing the recognition of an objective sleep epoch constitutes the context. In this case, the context is determined by the kernel size of the 1D convolutions that make up the second submodel of the CNN architecture. In the previous analysis, the kernel size was set to 3 to make it equivalent to the control scenario. Furthermore, we employed different sizes to mimic the way manual scoring is done by human experts and investigate the impact of temporal dynamics in offline decoding (see Table 4 and Supplementary Table S2). We can observe that looking only at the target epoch (kernel=1) produces a slight decrease in accuracy, while the highest performance is achieved with kernel sizes of 3 and 11 (past and future context available up to 30 seconds and 2.5 minutes, respectively). However, the model does not leverage relevant features appearing in longer temporal contexts. Therefore, for the proposed model, it seems unnecessary to handle large amounts of consecutive epochs around the one under recognition. The best compromise between performance and computational cost is achieved with the kernel of size 3, and the non-causal decoding methodology only outperforms the causal methodology by 2% (80.65% vs. 78.47%).

**Table 4:** Results overview of non-causal configurations. Accuracy achieved for each dataset in the control scenario and every kernel size for Experiment 4.

| Accuracy (%) | Control | Exp. 4<br>(kernel=1) | Exp. 4<br>(kernel=3) | Exp. 4<br>(kernel=11) | Exp. 4<br>(kernel=21) | Exp. 4<br>(kernel=41) |
| --- | --- | --- | --- | --- | --- | --- |
| SleepEDF | 80.97 | 77.58 | 82.03 | 81.70 | 80.81 | 79.45 |
| DOD-H | 81.92 | 78.79 | 82.91 | 83.25 | 77.49 | 75.63 |
| ISRUC | 71.69 | 72.84 | 76.27 | 76.33 | 71.82 | 62.01 |
| DOD-O | 79.31 | 76.54 | 81.38 | 79.78 | 78.17 | 76.16 |
| **Average** | 78.47 | 76.44 | 80.65 | 80.23 | 77.07 | 73.31 |

### Differences in sleep decoding between healthy participants vs. patients

The final analysis compares the performance of sleep scoring between patients and healthy participants. Given that this process is commonly used to identify and diagnose various disorders, understanding how automatic algorithms perform on fragmented sleep is crucial for clinical translation. To address this, two of the datasets we included consisted of patients suffering from sleep apnea (ISRUC, n = 100; and DOD-O, n = 50; see Figure 7A). Furthermore, we are specifically interested in understanding if there are notable differences between healthy subjects and patients when performing real-time single-channel decoding, as this modality holds the most potential to become part of take-home solutions for sleep disorders. A single channel allows for the most comfortable setup and real-time staging is a prerequisite for any type of closed-loop sleep improvement technique.

**Figure 7:**
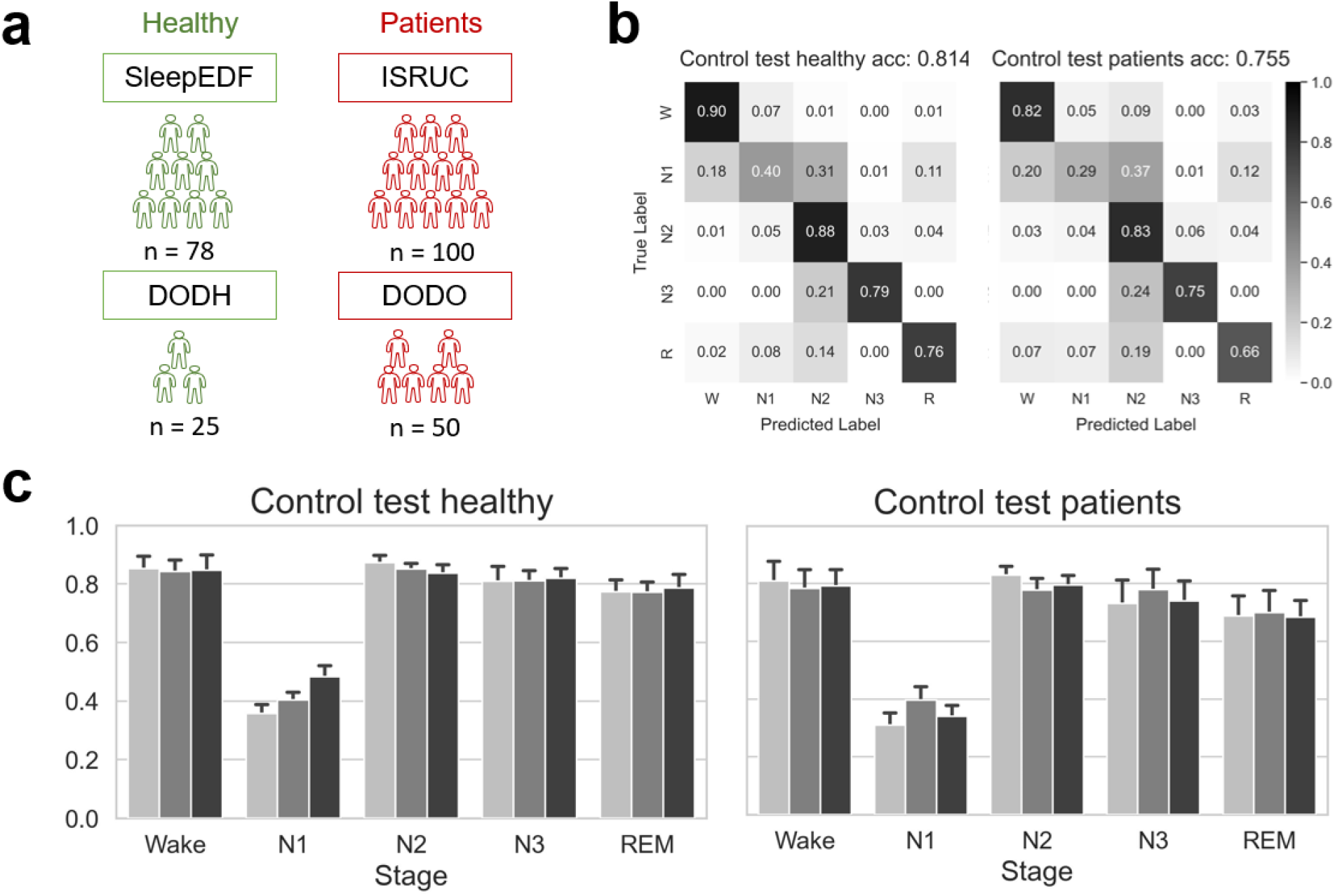
Comparison of decoding performance in recordings from individuals with and without sleep apnea. Panel A outlines the four open-access sleep datasets employed in the analysis. Two of them include recordings from healthy individuals (in green: SleepEDF, n=78; DOD-H, n=25), while the other two comprise subjects with sleep apnea (in red: ISRUC, n=100, DOD-O, n=50). Panel B displays the confusion matrices alongside the accuracy values obtained in the healthy and patient cohorts. Panel C contains bar plots displaying the values of recall (light grey), F1-score (dark grey), and precision (black).

Characteristic sleep fragmentation of patients with sleep apnea presents challenges in manual scoring. As expected, this complexity also affects automatic decoding, resulting in a drop in classification performance. Our results reveal an overall decrease in accuracy for the patient cohorts compared to the healthy cohorts (75.5% vs 81.4%, see Figure 7B). The drop in decoding accuracy per class is evident across all sleep phases, ranging from 4% in N3 to 11% in N1, and is similarly reflected in precision, recall and F1 score metrics (Figure 7C).

## Discussion

The global demographic shift towards an aging population highlights the pressing need for accessible and efficient solutions to address widespread healthcare challenges (Bardhan et al. 2020). Among these challenges, sleep disorders represent a significant public health concern affecting a substantial portion of the adult population (Patel et al. 2018; Pavlova and Latreille 2019). Digital therapeutics offer a promising avenue for tackling these issues, e.g., in the form of innovative solutions for sleep assessment and management (Bah et al. 2019). Automatic sleep scoring stands out as a pivotal component in this domain, facilitating timely diagnoses and personalized treatment plans.

This study aimed to explore various aspects crucial for the design of an automatic sleep scoring system, including electrode setup (number and type of sensors used), temporal scope (the inclusion or exclusion of future information to score an epoch), and the population sample (distinguishing between participants with and without sleep disorders). To conduct our analyses, we employed an existing CNN architecture based on 1D convolutions capable of analyzing sleep stages in real-time from raw single-channel EEG data or multi-channel multimodal data (Youness 2020). The maximum decoding accuracy results achieved with this model align with the current state of the art (see Table 1) and fall within the range of human inter-scorer variability (Danker-Hopfe et al. 2009; Rosenberg and Van Hout 2013).

Our investigation yielded three main contributions. Firstly, we found that sleep scoring based on a single frontal electrode demonstrated reliable congruency with human expert scorers. The network achieved an accuracy of 81.4% for datasets that included only healthy participants, and an average accuracy of 78.5% across all studied datasets, which also included individuals with sleep disorders. Interestingly, incorporating additional cortical locations or other physiological signals did not improve decoding accuracy. Secondly, utilizing future information to score a 30-second window did not lead to substantial improvements in decoding accuracy. Therefore, real-time sleep scoring can be accomplished with only a small decrease (around 2%) compared to purely offline methods that have access to the complete recording. Thirdly, we observed a notable reduction in decoding accuracy among individuals with sleep disorders compared to those without. Our analysis revealed an average drop of around 6% in accuracy for datasets that included patients. This finding emphasizes the challenges posed by sleep disorders and the growing importance of tailored approaches in sleep assessment and management.

### Relevance of EEG channels and additional sensors in real-time decoding

The impact of frontal EEG electrodes in sleep staging is particularly notable in our study, performing comparably to central locations and outperforming occipital ones. These results align with prior studies examining channel contributions in end-to-end staging approaches. For instance, (Chambon et al. 2018) noted the lowest accuracy with occipital electrodes, while (Lu et al. 2023) reported that central electrodes were the most informative, consistent with AASM scoring guidelines. Similarly, (Supratak et al. 2017) observed a decrease in staging performance when transitioning from fronto-central to parieto-occipital derivations. Frontal locations may simply be sufficient for capturing most stage-specific markers to approximate human scorer levels. For example, a major hallmark of non-REM sleep, sleep spindles, tend to spread at least to some extent to frontal sites. Slow waves, a marker for deep non-REM sleep, even predominates over frontal areas. Therefore, occipital locations might fail to capture transitions between specific phases like REM or N1, as reflected in our results.

Adding information from central and occipital channels to frontal ones did not significantly enhance overall accuracy in our experiments. Chambon et al. 2018 also explored additional sensors for automatic staging, noting a performance increase up to 6 electrodes. Similarly, (Biswal et al. 2018) found accuracy differences favoring 6-channel staging over 2-channel staging. Our study did not observe a similar effect, possibly due to differences in the network implementations or studied datasets. Note that our comparison was limited to 3 channels across 4 datasets. Future research could extend this analysis to include all possible channels across these and other datasets. Overall, our results confirm previous findings that fronto-central electrodes are optimal for sleep scoring, maintaining accuracy between offline and real-time configurations. Furthermore, we demonstrated that multiple EEG sensors do not enhance real-time staging accuracy in the proposed model.

### Temporal Scope and Real-Time Decoding

Our investigation into the temporal scope for automatic sleep scoring yielded a significant finding for real-time applications. The model achieved similar decoding accuracy when relying solely on past information (78.5% average across all datasets), compared to its best offline configuration (80.6%), which considered 30 seconds of future data to score an epoch. Notably, extending the temporal window to include longer future intervals did not enhance performance (80.2% for 2.5 minutes, 77.1% for 5 minutes, 73.3% for 10 minutes). This 2% difference between offline and real-time compatible approaches aligns with findings in the literature, as reflected in Table 1, where methods using only past data show similar differences compared to those incorporating future data.

The feasibility of accurate sleep scoring without the need for future information holds significant implications for digital therapeutic interventions. Currently, most applications for automatic sleep scoring operate offline, for instance to support diagnostics, or to analyze full-night recordings for research purposes. However, more and more studies are proving their feasibility for online applications, such as real-time sleep monitoring or closed-loop neurostimulation (Seo et al. 2020; Supratak and Guo 2020; Eldele et al. 2021; Fu et al. 2021). Therefore, trade-offs should be carefully considered, opting for either slightly superior decoding accuracy in offline settings or approaches that can be used in real-time with minimal performance loss.

### Potential Implications for Clinical Populations

The observed decrease in decoding accuracy for participants with sleep disorders compared to healthy participants is a crucial finding of this study. While the overall accuracy of the model with individuals without diagnosed sleep problems (81.4%) remains promising and aligns with most published results (cf. Table 1), the decline when evaluating patient data (75.5%) prompts further consideration for applications in clinical settings. This is particularly relevant given that sleep staging is fundamental for diagnosing and managing various sleep disorders.

Several factors may contribute to the observed decrease in accuracy for patients with sleep disorders, specifically sleep apnea in this study. The size of the datasets did not seem to play a role in this respect, as ISRUC and DOD-O datasets involve more participants than SleepEDF and DOD-H, respectively, and still provided lower decoding accuracy results. Sleep apnea is characterized by frequent arousals, disrupted sleep patterns (i.e., sleep fragmentation), and a higher prevalence of movement or other artifacts during sleep. These factors can negatively impact the quality of the EEG recordings and, consequently, the model’s performance. Variability in manual sleep scoring among these patients exists even for trained experts (Lee et al. 2020), underscoring the challenges inherent in accurately assessing sleep stages in clinical populations. In fact, the average accuracy we obtained with OSA patients (75.5%) is consistent with reported inter-rater agreement with this population (Danker-Hopfe et al. 2004).

The application of automatic sleep scoring in clinical settings, particularly for patients with sleep disorders, holds promising prospects. However, there is currently no well-defined threshold for clinically acceptable accuracy in automatic sleep scoring across different patient groups. Targeted optimizations may potentially enhance the models’ performance to levels achievable with healthy participants. Nevertheless, our findings emphasize the importance of further research on datasets encompassing a broader range of sleep disorders and severities to ensure that the model performs reliably across different patient profiles. Including the detection of disorder-associated events, such as apneas and hypopneas, and using them as features in a clinical sleep staging model, might help to overcome the described performance decrease on patient data.

### Applications and future directions

The potential applications of automatic sleep scoring extend far beyond research settings and hold significant promise for clinical practice. In clinics, sleep doctors could integrate automatic sleep scoring tools to streamline the diagnostic process and facilitate timely interventions. By automating the labor-intensive task of sleep staging, clinicians could allocate more time to patient care and decision-making, ultimately improving the quality and efficiency of healthcare delivery (Gaiduk et al. 2023; Soleimani et al. 2023). Moreover, the utility of automatic sleep scoring extends beyond the clinic and home-based monitoring may facilitate earlier detection of sleep disorders. Wearable devices equipped with a reduced number of sensors could be employed for data collection, making sleep assessments more accessible and convenient for patients (Liang and Chapa Martell 2018; Carneiro et al. 2020; López-Larraz et al. 2023; de Gans et al. 2024). While human experts may struggle with sleep scoring based on limited sensor data, algorithms such as the one presented here can still achieve high levels of accuracy.

Despite the potential benefits of this automatization, widespread adoption of these systems in clinical practice remains limited. One of the primary concerns lies in the opacity of black-box algorithms, which raises questions about their reliability and transparency (Loyola-Gonzalez 2019). Improving the explainability and interpretability of these methods will be crucial for their actual implantation, for instance, by investigating whether clinicians would make the same diagnostic and treatment decisions based on manual scorings versus those obtained from automated algorithms (Ratti and Graves 2022; Yang et al. (2022)).

Apart from its application in sleep monitoring, automatic sleep scoring can play a crucial role in developing and refining sleep interventions. Closed-loop neurostimulation has emerged as a promising approach for sleep disorders, with ongoing research demonstrating its potential (Ngo et al. 2013; Garcia-Molina et al. 2018; Esparza-Iaizzo et al. 2021). Automatic sleep scoring systems, like the one presented in this study, are an integral part of the functioning, validation, and optimization of most real time interventions. Ultimately, automatic sleep scoring has the potential to revolutionize the landscape of sleep medicine. As research progresses and neurostimulation interventions become more refined, these systems hold promise as invaluable clinical tools for optimizing sleep health and enhancing patient outcomes (Bah et al. 2019; Lee et al. 2020).

In conclusion, our study provides valuable insights into the boundary conditions of high-performance automatic sleep scoring and offers considerations for the development of effective and efficient sleep assessment tools. Still, it is important to note that a limitation of our study is that the results might be dependent on the chosen CNN architecture. Therefore, future research should validate if our findings also apply to other types of architectures, such as RNNs or attention-based models. Our results on the nuanced effects of electrode setup, temporal scope, and population characteristics will inform the work of all parties involved in the development and use of automatic sleep scoring systems, may it be engineers, researchers, or clinicians.

## Authors’ contributions

MEI, JM, LM and ELL conceived and designed the study. MEI and MST processed the data and obtained the results. MEI, MST, JGK, JM, LM and ELL analyzed and interpreted the results. MEI and ELL drafted the manuscript with inputs from MST and JGK. All authors revised and approved the manuscript.

## Supplementary

**Table S1:**
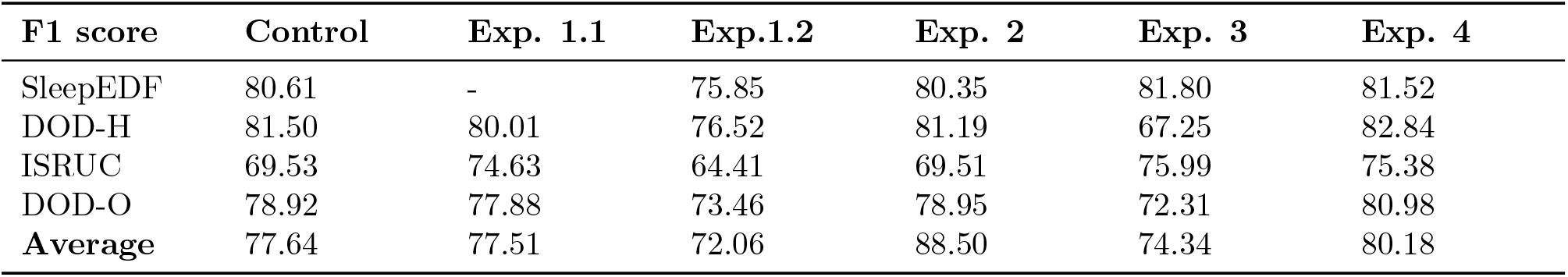
F1 score overview. Values of F1 score achieved for each dataset in the control scenario and in the different evaluation experiments.

**Table S2:** F1 score overview of non-causal configurations. Values of F1 score achieved for each dataset in the control scenario and every kernel size for Experiment 4.

| Accuracy (%) | Control | Exp. 1.1 | Exp.1.2 | Exp. 2 | Exp. 3 | Exp. 4 |
| --- | --- | --- | --- | --- | --- | --- |
| SleepEDF | 80.61 | 76.77 | 81.52 | 81.13 | 80.30 | 78.75 |
| DOD-H | 81.50 | 77.69 | 82.84 | 82.71 | 76.60 | 74.83 |
| ISRUC | 69.53 | 71.32 | 75.38 | 75.27 | 69.79 | 58.00 |
| DOD-O | 78.92 | 75.23 | 80.98 | 79.33 | 77.59 | 75.57 |
| **Average** | 77.64 | 75.25 | 80.18 | 79.61 | 76.07 | 71.79 |

## References

Abadi, M., Barham, P., Chen, J., Chen, Z., Davis, A., Dean, J., et al. (2016). TensorFlow: A System for Large-Scale Machine Learning. In Proceedings of the 12th USENIX Symposium on Operating Systems Design and Implementation (OSDI ‘16). 265–283

Abbadessa, G., Brigo, F., Clerico, M., De Mercanti, S., Trojsi, F., Tedeschi, G., et al. (2022). Digital therapeutics in neurology. Journal of Neurology 269, 1209–1224. doi:10.1007/s00415-021-10608-4

Andreotti, F., Phan, H., Cooray, N., Lo, C., Hu, M. T. M., and De Vos, M. (2018). Multichannel Sleep Stage Classification and Transfer Learning using Convolutional Neural Networks. In 40th Annual International Conference of the IEEE Engineering in Medicine and Biology Society (EMBC) (IEEE), 171–174. doi:10.1109/EMBC.2018.8512214

Bah, T. M., Goodman, J., and Iliff, J. J. (2019). Sleep as a Therapeutic Target in the Aging Brain. Neurotherapeutics 16, 554–568. doi:10.1007/s13311-019-00769-6

Bardhan, I., Chen, H., and Karahanna, E. (2020). Connecting systems, data, and people: A multidisciplinary research roadmap for chronic disease management. MIS Quarterly: Management Information Systems 44, 185–200. doi:10.25300/MISQ/2020/14644

Berry, R. B., Brooks, R., Gamaldo, C. E., Harding, S. M., Lloyd, R. M., Marcus, C. L., et al. (2015). The AASM Manual for the Scoring of Sleep and Associated Events: Rules, Terminology and Technical Specifications, Version 2.2. Tech. rep., American Academy of Sleep Medicine, Darien, Illinois

Biswal, S., Sun, H., Goparaju, B., Westover, M. B., Sun, J., and Bianchi, M. T. (2018). Expert-level sleep scoring with deep neural networks. Journal of the American Medical Informatics Association 25, 1643–1650. doi:10.1093/jamia/ocy131

Brandmayr, G., Hartmann, M., Fürbass, F., Matz, G., Samwald, M., Kluge, T., et al. (2022). Relational local electroencephalography representations for sleep scoring. Neural Networks 154, 310–322. doi:10.1016/J.NEUNET.2022.07.020

Carneiro, M. R., de Almeida, A. T., and Tavakoli, M. (2020). Wearable and Comfortable e-Textile Headband for Long-Term Acquisition of Forehead EEG Signals. IEEE Sensors Journal 20, 15107–15116. doi:10.1109/JSEN.2020.3009629

Chambon, S., Galtier, M. N., Arnal, P. J., Wainrib, G., and Gramfort, A. (2018). A Deep Learning Architecture for Temporal Sleep Stage Classification Using Multivariate and Multimodal Time Series. IEEE Transactions on Neural Systems and Rehabilitation Engineering 26, 758–769. doi:10.1109/TNSRE.2018.2813138

[Dataset] Chollet, F. (2018). Keras: The Python Deep Learning library

Coon, W. G. and Ogg, M. (2024). Laying the Foundation: Modern Transformers for Gold-Standard Sleep Analysis. bioRxiv doi:10.1101/2024.01.18.576246

Danker-Hopfe, H., Anderer, P., Zeitlhofer, J., Boeck, M., Dorn, H., Gruber, G., et al. (2009). Interrater reliability for sleep scoring according to the Rechtschaffen Kales and the new AASM standard. Journal of Sleep Research 18, 74–84. doi:10.1111/j.1365-2869.2008.00700.x

Danker-Hopfe, H., Kunz, D., Gruber, G., Klösch, G., Lorenzo, J. L., Himanen, S. L., et al. (2004). Interrater reliability between scorers from eight European sleep laboratories in subjects with different sleep disorders. Journal of Sleep Research 13, 63–69. doi:10.1046/j.1365-2869.2003.00375.x

de Gans, C., Burger, P., Van den Ende, E., Hermanides, J., Nanayakkara, P., Gemke, R., et al. (2024). Sleep assessment using EEG-based wearables – A systematic review. Sleep Medicine Reviews, 101951 doi:10.1016/J.SMRV.2024.101951

Dutt, M., Redhu, S., Goodwin, M., and Omlin, C. W. (2023). SleepXAI: An explainable deep learning approach for multi-class sleep stage identification. Applied Intelligence 53, 16830–16843. doi:10.1007/s10489-022-04357-8

Eldele, E., Chen, Z., Liu, C., Wu, M., Kwoh, C.-K., Li, X., et al. (2021). An Attention-Based Deep Learning Approach for Sleep Stage Classification With Single-Channel EEG. IEEE Transactions on Neural Systems and Rehabilitation Engineering 29, 809–818. doi:10.1109/TNSRE.2021.3076234

Escourrou, P., Luriau, S., Rehel, M., Nédelcoux, H., and Lanöe, J.-L. (2000). Needs and Costs of Sleep Monitoring. Studies in Health Technology and Informatics 78, 69–85. doi:10.3233/978-1-60750-922-6-69

Esparza-Iaizzo, M., Álvarez, I., Klinzing, J. G., Montesano, L., Minguez, J., and López-Larraz, E. (2021). SleepBCI: a platform for memory enhancement during sleep based on automatic scoring. In XXXIX Annual Congress of the Spanish Society of Biomedical Engineering (Valladolid)

Fu, G., Zhou, Y., Gong, P., Wang, P., Shao, W., and Zhang, D. (2023). A Temporal-Spectral Fused and Attention-Based Deep Model for Automatic Sleep Staging. IEEE Transactions on Neural Systems and Rehabilitation Engineering 31, 1008–1018. doi:10.1109/TNSRE.2023.3238852

Fu, M., Wang, Y., Chen, Z., Li, J., Xu, F., Liu, X., et al. (2021). Deep Learning in Automatic Sleep Staging With a Single Channel Electroencephalography. Frontiers in Physiology 12, 628502. doi:10.3389/fphys.2021.628502

Gaiduk, M., Serrano Alarcón, Á., Seepold, R., and Martínez Madrid, N. (2023). Current status and prospects of automatic sleep stages scoring: Review. Biomedical Engineering Letters 13, 247–272. doi:10.1007/s13534-023-00299-3

Garcia-Molina, G., Tsoneva, T., Jasko, J., Steele, B., Aquino, A., Baher, K., et al. (2018). Closedloop system to enhance slow-wave activity. Journal of Neural Engineering 15, 066018. doi:10.1088/1741-2552/aae18f

Goldberger, A. L., Amaral, L. A. N., Glass, L., Hausdorff, J. M., Ivanov, P. C., Mark, R. G., et al. (2000). PhysioBank, PhysioToolkit, and PhysioNet: Components of a New Research Resource for Complex Physiologic Signals. Circulation 101, e215–e220. doi:10.1161/01.CIR.101.23.e215

Guillot, A., Sauvet, F., During, E. H., and Thorey, V. (2020). Dreem Open Datasets: Multi-Scored Sleep Datasets to Compare Human and Automated Sleep Staging. IEEE Transactions on Neural Systems and Rehabilitation Engineering 28, 1955–1965. doi:10.1109/TNSRE.2020.3011181

Imtiaz, S. A. and Rodriguez-Villegas, E. (2014). A Low Computational Cost Algorithm for REM Sleep Detection Using Single Channel EEG. Annals of Biomedical Engineering 42, 2344–2359. doi:10.1007/s10439-014-1085-6

Kemp, B., Zwinderman, A., Tuk, B., Kamphuisen, H., and Oberye, J. (2000). Analysis of a sleep-dependent neuronal feedback loop: the slow-wave microcontinuity of the EEG. IEEE Transactions on Biomedical Engineering 47, 1185–1194. doi:10.1109/10.867928

Khalighi, S., Sousa, T., Santos, J. M., and Nunes, U. (2016). ISRUC-Sleep: A comprehensive public dataset for sleep researchers. Computer Methods and Programs in Biomedicine 124, 180–192. doi: 10.1016/j.cmpb.2015.10.013

Kingma, D. P. and Ba, J. (2014). Adam: A Method for Stochastic Optimization. arXiv preprint arXiv, 1412.6980. doi:10.48550/arXiv.1412.6980

Lee, S., Yu, Y., Back, S., Seo, H., and Lee, K. (2024). SleePyCo: Automatic sleep scoring with feature pyramid and contrastive learning. Expert Systems with Applications 240, 122551. doi: 10.1016/J.ESWA.2023.122551

Lee, Y. F., Gerashchenko, D., Timofeev, I., Bacskai, B. J., and Kastanenka, K. V. (2020). Slow Wave Sleep Is a Promising Intervention Target for Alzheimer’s Disease. Frontiers in Neuroscience 14, 705. doi:10.3389/fnins.2020.00705

Lemkhenter, A. and Favaro, P. (2022). Towards Sleep Scoring Generalization Through Self-Supervised Meta-Learning. In 44th Annual International Conference of the IEEE Engineering in Medicine Biology Society (EMBC). 2961–2966. doi:10.1109/EMBC48229.2022.9871056

Liang, Z. and Chapa Martell, M. A. (2018). Validity of Consumer Activity Wristbands and Wearable EEG for Measuring Overall Sleep Parameters and Sleep Structure in Free-Living Conditions. Journal of Healthcare Informatics Research 2, 152–178. doi:10.1007/s41666-018-0013-1

López-Larraz, E., Escolano, C., Robledo-Menéndez, A., Morlas, L., Alda, A., and Minguez, J. (2023). A garment that measures brain activity: proof of concept of an EEG sensor layer fully implemented with smart textiles. Frontiers in Human Neuroscience 17, 1135153. doi: 10.3389/fnhum.2023.1135153

Loyola-Gonzalez, O. (2019). Black-Box vs. White-Box: Understanding Their Advantages and Weak-nesses From a Practical Point of View. IEEE Access 7, 154096–154113. doi:10.1109/ACCESS.2019.2949286

Lu, C., Pathak, S., Englebienne, G., and Seifert, C. (2023). Channel Contribution in Deep Learning Based Automatic Sleep Scoring—How Many Channels Do We Need? IEEE Transactions on Neural Systems and Rehabilitation Engineering 31, 494–505. doi:10.1109/TNSRE.2022.3227040

Makin, S. (2019). A smarter way to treat. Nature 573, S106–S109. doi:10.1038/nrendo.2017.135

Mousavi, S., Afghah, F., and Acharya, U. R. (2019). SleepEEGNet: Automated sleep stage scoring with sequence to sequence deep learning approach. PLoS ONE 14, e0216456. doi:10.1371/journal.pone.0216456

Ngo, H.-V., Martinetz, T., Born, J., and Mölle, M. (2013). Auditory Closed-Loop Stimulation of the Sleep Slow Oscillation Enhances Memory. Neuron 78, 545–553. doi:10.1016/J.NEURON.2013.03.006

O’Reilly, C., Gosselin, N., Carrier, J., and Nielsen, T. (2014). Montreal Archive of Sleep Studies: an open-access resource for instrument benchmarking and exploratory research. Journal of Sleep Research 23, 628–635. doi:10.1111/jsr.12169

Park, K. M., Lee, S., and Lee, E. (2021). Can Digital Therapeutics Open a New Era of Sleep Medicine? Chronobiology in Medicine 3, 142–148. doi:10.33069/CIM.2021.0028

Patel, D., Steinberg, J., and Patel, P. (2018). Insomnia in the Elderly: A Review. Journal of Clinical Sleep Medicine 14, 1017–1024. doi:10.5664/jcsm.7172

Pavlova, M. K. and Latreille, V. (2019). Sleep Disorders. The American Journal of Medicine 132, 292–299. doi:10.1016/j.amjmed.2018.09.021

Perslev, M., Darkner, S., Kempfner, L., Nikolic, M., Jennum, P. J., and Igel, C. (2021). U-Sleep: resilient high-frequency sleep staging. npj Digital Medicine 4, 72. doi:10.1038/s41746-021-00440-5

Perslev, M., Jensen, M. H., Darkner, S., Jennum, P. J., and Igel, C. (2019). U-Time: A Fully Convolutional Network for Time Series Segmentation Applied to Sleep Staging. In Proceedings of the 33rd International Conference on Neural Information Processing Systems (NIPS). 4415–4426

Phan, H., Andreotti, F., Cooray, N., Chén, O. Y., and De Vos, M. (2019). SeqSleepNet: End-to-End Hierarchical Recurrent Neural Network for Sequence-to-Sequence Automatic Sleep Staging. IEEE Transactions on Neural Systems and Rehabilitation Engineering 27, 400–410. doi:10.1109/TNSRE.2019.2896659

Phan, H., Chén, O. Y., Tran, M. C., Koch, P., Mertins, A., and De Vos, M. (2021). XSleepNet: Multi-View Sequential Model for Automatic Sleep Staging. IEEE Transactions on Pattern Analysis and Machine Intelligence 44, 5903–5915. doi:10.1109/TPAMI.2021.3070057

Phan, H., Lorenzen, K. P., Heremans, E., Chén, O. Y., Chén, C., Tran, M. C., et al. (2023). L-SeqSleepNet: Whole-cycle Long Sequence Modelling for Automatic Sleep Staging. IEEE Journal of Biomedical and Health Informatics 27, 4748–4757. doi:10.1109/JBHI.2023.3303197

Phan, H., Mikkelsen, K., Chen, O. Y., Koch, P., Mertins, A., and De Vos, M. (2022). SleepTransformer: Automatic Sleep Staging With Interpretability and Uncertainty Quantification. IEEE Transactions on Biomedical Engineering 69, 2456–2467. doi:10.1109/TBME.2022.3147187

Pradeepkumar, J., Anandakumar, M., Kugathasan, V., Suntharalingham, D., Kappel, S. L., De Silva, A. C., et al. (2022). Towards Interpretable Sleep Stage Classification Using Cross-Modal Transformers. arXiv preprint 2208, 06991. doi:10.48550/arXiv.2208.06991

Quan, S. F., Howard, B. V., Iber, C., Kiley, J. P., Nieto, F. J., O’Connor, G. T., et al. (1997). The Sleep Heart Health Study: Design, Rationale, and Methods. Sleep 20, 1077–1085. doi:10.1093/sleep/20.12.1077

Ratti, E. and Graves, M. (2022). Explainable machine learning practices: opening another black box for reliable medical AI. AI and Ethics 2, 801–814. doi:10.1007/s43681-022-00141-z

Rosenberg, R. S. and Van Hout, S. (2013). The American Academy of Sleep Medicine inter-scorer reliability program: sleep stage scoring. Journal of clinical sleep medicine : JCSM : official publication of the American Academy of Sleep Medicine 9, 81–7. doi:10.5664/jcsm.2350

Schaltenbrand, N., Lengelle, R., Toussaint, M., Luthringer, R., Carelli, G., Jacqmin, A., et al. (1996). Sleep Stage Scoring Using the Neural Network Model: Comparison Between Visual and Automatic Analysis in Normal Subjects and Patients. Sleep 19, 26–35. doi:10.1093/sleep/19.1.26

Seo, H., Back, S., Lee, S., Park, D., Kim, T., and Lee, K. (2020). Intra- and inter-epoch temporal context network (IITNet) using sub-epoch features for automatic sleep scoring on raw single-channel EEG. Biomedical Signal Processing and Control 61, 102037. doi:10.1016/J.BSPC.2020.102037

Sharma, S. K., Maiti, S., Mythirayee, S., Srijithesh, P. R., and Bapi, R. S. (2023). Transparency in Sleep Staging: Deep Learning Method for EEG Sleep Stage Classification with Model Interpretability. arXiv preprint 2309, 07156v4. doi:10.48550/arXiv.2309.07156

Sivertsen, B., Lallukka, T., and Salo, P. (2011). The Economic Burden of Insomnia at the Workplace. An Opportunity and Time for Intervention? Sleep 34, 1151–1152. doi:10.5665/SLEEP.1224

Soleimani, R., Barahona, J., Chen, Y., Bozkurt, A., Daniele, M., Pozdin, V., et al. (2023). Advances in Modeling and Interpretability of Deep Neural Sleep Staging: A Systematic Review. Physiologia 4, 1–42. doi:10.3390/physiologia4010001

Supratak, A., Dong, H., Wu, C., and Guo, Y. (2017). DeepSleepNet: A model for automatic sleep stage scoring based on raw single-channel EEG. IEEE Transactions on Neural Systems and Rehabilitation Engineering 25, 1998–2008. doi:10.1109/TNSRE.2017.2721116

Supratak, A. and Guo, Y. (2020). TinySleepNet: An Efficient Deep Learning Model for Sleep Stage Scoring based on Raw Single-Channel EEG. In 42nd Annual International Conference of the IEEE Engineering in Medicine and Biology Society (EMBC). 641–644. doi:10.1109/EMBC44109.2020.9176741

Tarasiuk, A. and Reuveni, H. (2013). The economic impact of obstructive sleep apnea. Current Opinion in Pulmonary Medicine 19, 639–644. doi:10.1097/MCP.0b013e3283659e1e

Van Rossum, G. and Drake, F. L. (1995). Python reference manual (Amsterdam: Centrum voor Wiskunde en Informatica)

Van Ryswyk, E. M., Mukherjee, S., Chai-Coetzer, C. L., Vakulin, A., and McEvoy, R. D. (2018). Sleep Disorders, Including Sleep Apnea, and Hypertension. American Journal of Hypertension 31, 857–864. doi:10.1093/ajh/hpy082

Yan, R., Li, F., Zhou, D. D., Ristaniemi, T., and Cong, F. (2021). Automatic sleep scoring: A deep learning architecture for multi-modality time series. Journal of Neuroscience Methods 348, 108971. doi:10.1016/j.jneumeth.2020.108971

Yang, G., Ye, Q., and Xia, J. (2022). Unbox the black-box for the medical explainable AI via multimodal and multi-centre data fusion: A mini-review, two showcases and beyond. Information Fusion 77, 29–52. doi:10.1016/J.INFFUS.2021.07.016

[Dataset] Youness, M. (2020). CVxTz/EEG classification: v1.0. doi:10.5281/zenodo.4060151

Zhang, Y., Zhang, X., Liu, W., Luo, Y., Yu, E., Zou, K., et al. (2014). Automatic Sleep Staging using Multi-dimensional Feature Extraction and Multi-kernel Fuzzy Support Vector Machine. Journal of Healthcare Engineering 5, 505–520. doi:10.1260/2040-2295.5.4.505

Zhou, W., Zhu, H., Shen, N., Chen, H., Fu, C., Yu, H., et al. (2023). A Lightweight Segmented Attention Network for Sleep Staging by Fusing Local Characteristics and Adjacent Information. IEEE Transactions on Neural Systems and Rehabilitation Engineering 31, 1008–1018. doi:10.1109/TNSRE.2022.3220372

